# Capturing G protein-coupled receptors into native lipid-bilayer nanodiscs using new diisobutylene/maleic acid (DIBMA) copolymers

**DOI:** 10.1101/2024.01.20.576420

**Authors:** Ci Chu, Carolyn Vargas, Maria Carolina Barbosa, Simon Sommerhage, Gunnar F. Schröder, Sandro Keller, Manuel Etzkorn

## Abstract

Many membrane proteins, including G-protein-coupled receptors (GPCRs), are susceptible to denaturation when extracted from their native membrane by detergents. Therefore, alternative methods have been developed, including amphiphilic copolymers that enable the direct extraction of functional membrane proteins along with their surrounding lipids, leading to the formation of native lipid-bilayer nanodiscs. Among these amphiphilic copolymers, styrene/maleic acid (SMA) and diisobutylene/maleic acid (DIBMA) polymers have been extensively studied and successfully utilized to isolate various types of membrane proteins, including GPCRs. Despite their many benefits, SMA and DIBMA polymers also have significant drawbacks that limit their application. Most notably, both SMA and DIBMA carry high negative charge densities, which can interfere with protein–protein and protein–lipid interactions through unspecific Coulombic attraction or repulsion. Herein, we describe a series of new amphiphilic copolymers derived from DIBMA via partial amidation of the carboxylate pendant groups with various biocompatible amines. The nanodisc-forming properties of the new polymers were assessed using model membranes as well as in the context of extracting the melanocortin 4 receptor (MC4R), a prototypical class A GPCR. While each new DIBMA variant displays distinct features that may be favorable for selected applications, we identified a new PEGylated DIBMA variant called mPEG_4_-DIBMA as particularly promising for the studied purpose. On the one hand, mPEG_4_-DIBMA abolishes unspecific interactions with the tested peptide ligand, a prerequisite for reliably characterizing GPCR–ligand interactions. On the other hand, mPEG_4_-DIBMA outperforms other polymers such as SMA and DIBMA by achieving higher extraction efficiencies of MC4R from Sf9 insect cell membranes. Thus, this new nanodisc-forming polymer combines two key advantages that are crucial for investigating GPCRs in a well-defined but still native lipid-bilayer environment, thus paving the way for manifold future applications.

## Introduction

G protein-coupled receptors (GPCRs), the largest superfamily of cell-surface proteins, share a conserved architecture of seven transmembrane helical domains (TMDs) and have been implicated in a plethora of diseases such as cancer, obesity, and Alzheimer’s disease.^1–3^ The isolation of GPCRs from cellular membranes for subsequent *in vitro* studies is traditionally carried out with the aid of detergents, which displace native lipids and form micelles as a membrane-mimetic environment to solubilize GPCRs. Lipid-bilayer nanodiscs formed by membrane scaffold proteins (MSPs) have also been widely used in studies of GPCRs,^4^ such as neurotensin receptor 1 (NTSR1),^5^ and rhodopsin.^6^ However, the stability of GPCRs often decreases upon solubilization by detergents, which is the first step for preparing MSP nanodiscs.^4^ In addition, GPCR activities are regulated by various physical properties of the surrounding phospholipid bilayer such as lipid order, lateral pressure, bilayer thickness, hydrophobic mismatch, membrane fluidity, curvature, and lipid composition, which are altered in detergent micelles and difficult to mimic in synthetic lipid mixtures.^7–10^ The lipid environment also plays critical roles in GPCR–ligand interactions, receptor coupling, and the recruitment of GPCR kinases (GRKs) and arrestins.^11–14^ Therefore, the study of GPCRs in more native-like environments is highly desirable.

In addition to model membrane systems,^15^ significant progress has been made in developing and exploiting lipid-bilayer nanodiscs encapsulated by amphiphilic copolymers that directly extract membrane proteins together with their surrounding lipids from cellular membranes.^16^ These native nanodiscs often preserve the structural and functional integrity of extracted proteins. One of the most commonly used copolymers is styrene/maleic acid polymer SMA(2:1), a negatively charged random polymer employed to purify membrane proteins from different cell types.^17–22^ However, the efficiency of membrane-protein extraction by SMA(2:1) and the characterization of SMA-encapsulated proteins are restricted by a rather narrow buffer range, excluding lower pH values and even low concentrations of divalent cations such as Mg^2+^ and Ca^2+^.^23,24^ New strategies could overcome some of the limitations in terms of buffer compatibility,^25,26^ but the quantification of encapsulated proteins and several other biophysical experiments are hampered by SMA’s strong absorption in the UV range, which is due to its aromatic styrene moieties. Diisobutylene/maleic acid (DIBMA) is an alternating copolymer that does not contain any aromatic groups but retains the ability to solubilize membrane proteins and lipids.^27^ Thus, one of the significant advantages of DIBMA is its lower absorption in the UV range. Moreover, DIBMA exhibits only a gentle impact on the lipid acyl chain order and a high resistance against divalent cations.^28^ DIBMA has been successfully used to extract a broad range of membrane proteins from different host cells, including rhomboid proteases,^29^ the membrane tether protein ZipA,^30^ the ATP Binding Cassette (ABC) transporter BmrA,^30^ the GPCRs adenosine A_2A_ receptor (A_2A_R)^30^ and calcitonin gene-related peptide receptor (CGRP),^30^ as well as the mechanosensitive channel of small conductance (MscS)-like channel YnaI.^31^ Nonetheless, the high charge densities carried by both SMA and DIBMA, which are due to their carboxylic acid pendant groups, can interfere with binding measurements using charged ligands through unspecific Coulombic interactions.

In this work, we describe a series of new amphiphilic copolymers, including Dab-DIBMA, Arg-DIBMA, Meg-DIBMA, and mPEG_4_-DIBMA obtained from DIBMA via partial amidation with various biocompatible amines. The formation of nanodiscs by exposing large unilamellar vesicles (LUVs) to polymers, the morphology of the resulting nanodiscs, and the thermotropic phase behavior of the encapsulated lipid bilayers were examined with the aid of dynamic light scattering (DLS), transmission electron microscopy (TEM), and differential scanning calorimetry (DSC), respectively. Additionally, microfluidic diffusional sizing (MDS) was used to gauge potential unspecific interactions between peptide ligands and polymers. The efficacies of the new polymers in extracting a prototypical GPCR were assessed by using the human melanocortin 4 receptor (MC4R) expressed in insect cells. The results of our study demonstrate that all of the polymers examined are able to form lipid-bilayer nanodiscs with narrow size distributions. Furthermore, all polymers can extract MC4R into native nanodiscs, providing new tools for the structural and functional characterization of GPRCs. Notably, we found that mPEG_4_-DIBMA does not display any unspecific interactions with a cationic peptide ligand and exhibits only low UV absorption. In addition, mPEG_4_-DIBMA is highly water-soluble, readily solubilizes phospholipid vesicles, and efficiently extracts MC4R from insect membranes. Taken together, our study demonstrates that mPEG_4_-DIBMA is an outstanding amphiphilic copolymer for investigating integral membrane proteins in their native lipid environment.

## Results and discussion

### Design and synthesis of copolymers

The goal of our synthetic approach was to reduce the charge density on the polymer backbone by derivatizing DIBMA through amidation of one of the two carboxylic acid groups in each repeating unit (Scheme 1). We have previously used the same strategy to produce Glyco-DIBMA, which offers several advantages over unmodified DIBMA, including a higher protein-extraction efficiency.^32,33^ Here, we sought to explore a larger chemical space through amidation using a set of structurally diverse amines, including: α-amino acids such as L-arginine (Arg) and L-2,4-diaminobutyric acid (Dab); the hexosamine meglumine (Meg); and the oligo (ethylene oxide) tetraethyleneglycol monomethyl ether amine (mPEG_4_). These compounds have in common that they contain at least one reactive amino group, are highly water-soluble, biocompatible, and net neutral once coupled to the polymer backbone through an amide linkage. We considered these properties important to reduce the charge density on the DIBMA polymer backbone while retaining its excellent water solubility. Four polymers were thus synthesized, henceforth referred to as Arg-DIBMA, Dab-DIBMA, Meg-DIBMA, and mPEG_4_-DIBMA.

**Scheme 1.**
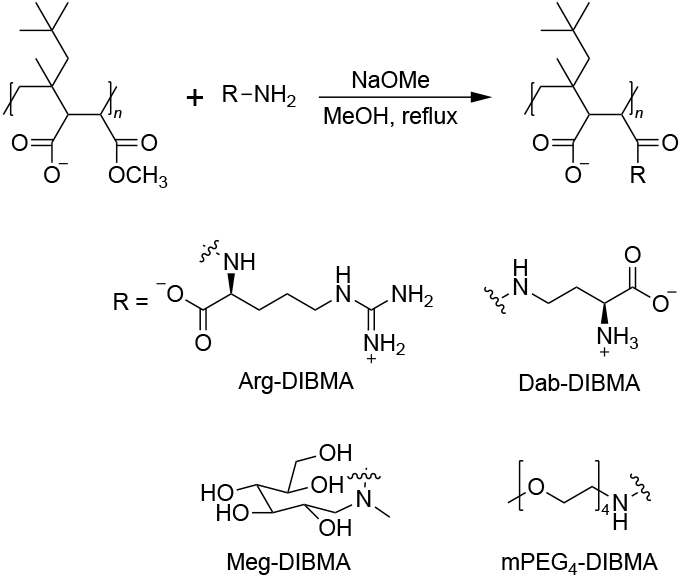
General scheme for the synthesis of DIBMA-based amphiphilic copolymers that form lipid-bilayer nanodiscs.

### Characterization of nanodisc-forming properties

Each polymer variant was subsequently tested for its key biophysical and nanodiscs-forming properties, including lipid extraction from synthetic vesicles as well as protein extraction from eukaryotic cells. First, we investigated the formation of polymer/DMPC nanodiscs and the solubilization efficiency of the new polymers. To this end, large unilamellar vesicles (LUVs) made from DMPC were incubated with each polymer at different polymer/lipid mass ratios, and particle size distributions were analyzed by dynamic light scattering (DLS). At a polymer/lipid mass ratio (*m*_P_*/m*_L_) of 2.0, the hydrodynamic particle sizes of all new nanodiscs were in the expected range of ∼10 nm (Fig. 1a). By contrast, at a lower *m*_P_*/m*_L_ of 0.5, substantial differences in hydrodynamic particle sizes were observed, reflecting differences in the solubilization efficiency among the polymers (Fig. 1b). A more systematic screening of *m*_P_*/m*_L_ revealed that mass ratios of ∼1 should be used for all new DIBMA variants to obtain the often-desired nanodisc diameter of ∼10 nm (Fig. 1c, top panel). This minimal *m*_P_*/m*_L_ ratio is considerably less than that observed for unmodified DIBMA but about a factor of two larger than for SMA(2:1) and Sulfo-DIBMA (Fig. 1c, lower panel). Sulfo-DIBMA is a recent electroneutral DIBMA variant that contains an imide ring,^34^ in contrast with the open amide linkage of the DIBMA variants reported here (Scheme 1).

**Figure 1.**
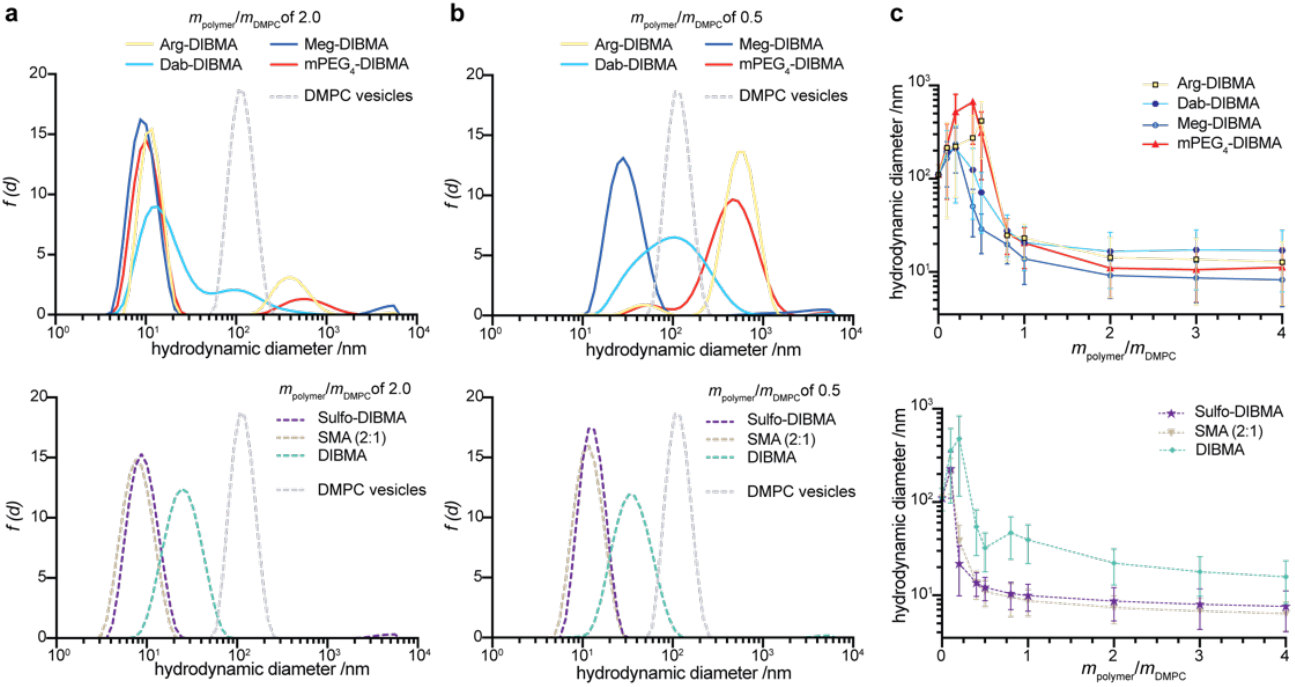
Particle size analysis shows formation of homogenous nanodiscs for all polymers at polymer/lipid mass ratios >1. (a,b) Intensity-weighted particle size distributions of mixtures containing polymer and DMPC (4 mg/mL) at polymer/lipid mass ratios (mP/mL) of (a) 2.0 and (b) 0.5 as obtained from DLS. Upper panels show data for the new DIBMA variants, while lower panels serve as reference for established polymers. (c) z-Average particle diameters derived from DLS as functions of mP/mL. Vertical bars indicate size distribution widths defined as σ = z * √PDI, where PDI is the polydispersity index.

### Suitability for interaction studies

Next, we tested the suitability of each new polymer for characterizing interactions that are susceptible to charge effects. Most existing polymers, such as SMA and DIBMA, are incompatible with ligand binding assays for various GPCRs because of unspecific interactions between the polyanionic polymer chains and the ligands.^34^ To assess the new DIBMA variants for their suitability in such ligand binding studies, we performed microfluidic diffusional sizing (MDS) measurements with the binding fragment of the adrenocorticotropic hormone (ACTH), a potent agonist of the melanocortin 4 receptor (MC4R).^35^ MDS allows for the determination of the hydrodynamic particle size of bare ACTH and its increase resulting from unspecific interactions with nanodiscs.^36,37^ To evaluate the interactions between the DIBMA variants and ACTH, we used polymer-encapsulated nanodiscs containing the zwitterionic phospholipid DMPC, a lipid known not to interact with ACTH.^34^ Polymer/DMPC nanodiscs were prepared at *m*_P_*/m*_L_ of 4 to obtain small nanodiscs, as confirmed by DLS (Fig. 1c and S1). In line with previous observations,^34^ our MDS results confirmed that the hydrodynamic particle size of ACTH remained unchanged upon addition of DMPC nanodiscs encapsulated by the electroneutral polymer Sulfo-DIBMA, indicating the absence of unspecific interactions between the polymer and the ligand (Fig. 2). In contrast with Sulfo-DIBMA, we observed an increase in hydrodynamic particle size when ACTH was exposed to nanodiscs encapsulated by DIBMA, Arg-DIBMA, Dab-DIBMA, or Meg-DIBMA, suggesting unspecific interactions between the cationic peptide ligand and the anionic polymers. In stark contrast, the hydrodynamic particle size of ACTH remained unchanged upon addition of nanodiscs encapsulated by the PEGylated polymer mPEG_4_-DIBMA, even though mPEG_4_-DIBMA still carries carboxylate groups (Scheme 1). For all polymers, the same interaction behaviors were found at two different pH values (pH 7.4, Fig. 2 and pH 8.0, Fig. S2), confirming that mPEG_4_-DIBMA is of particular interest for studying the specific binding of ligands to nanodiscs-embedded membrane proteins under commonly applied buffer conditions.

**Figure 2.**
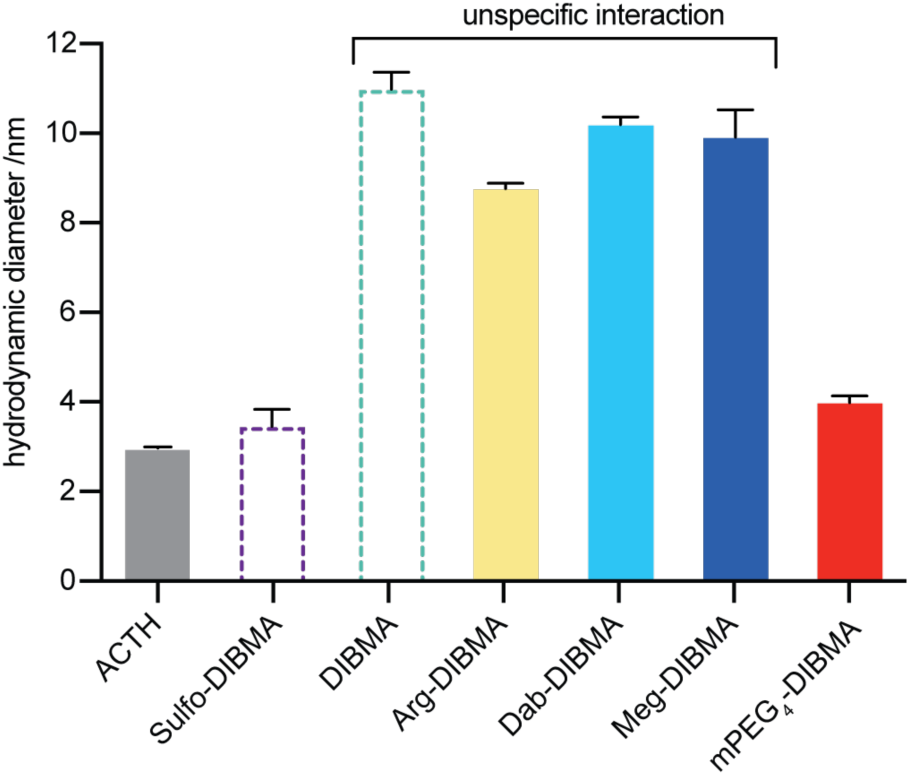
Only mPEG_4_-DIBMA shows reduced unspecific interactions with the cationic peptide ligand ACTH. Unspecific interactions between ACTH and polymer/DMPC nanodiscs at m_P_/m_L_ of 4 were measured by means of microfluidic diffusional sizing (MDS). All experiments were carried out at 50 mM HEPES, 200 mM NaCl, pH 7.4. Error bars represent one standard deviation of two independent experiments, each repeated in triplicate.

### Physicochemical properties of mPEG_**4**_**-DIBMA**

Due to its promising features in ligand-binding assays, we carried out a more comprehensive characterization of mPEG_4_-DIBMA. First, we investigated the origin of the reduced unspecific interactions with ACTH by determining the *ζ*-potential. Our results confirmed that unmodified DIBMA indeed has a strongly negative *ζ*-potential, which is only partly reduced by the mPEG_4_ modification, in contrast with electroneutral Sulfo-DIBMA (Fig. 3a). Negative *ζ*-potentials were similarly detected in lipid-free polymer samples and for lipid-encapsulating nanodiscs. From this finding, we infer that unspecific binding to mPEG_4_-DIBMA is not prevented by a lack of charged groups on the polymer but rather by the relatively bulky, strongly hydrated PEG chains, which sterically block access of the peptide to the polymer backbone. This interpretation is consistent with the expected Debye–Hückel screening length of <1 nm under the applied conditions, suggesting, that the unspecific interactions between the positively charged peptide and the negatively charged carboxylate groups are much less prominent in mPEG_4_-DIBMA because of their spatial separation by the hydrated PEG chains and the shielding effects of the buffer ions.

**Figure 3.**
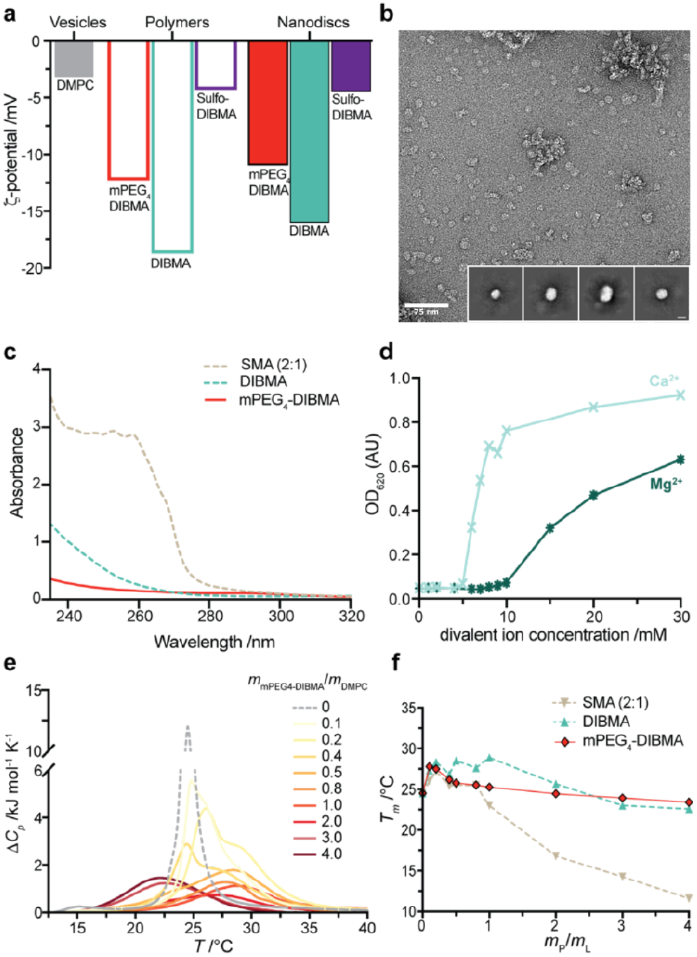
Physicochemical properties of mPEG_4_-DIBMA. (a) ζ-potentials of mPEG_4_-DIBMA, DIBMA, Sulfo-DIBMA, and respective polymer-encapsulated DMPC nanodiscs. (b) Negative-stain transmission electron microscopy (TEM) image of mPEG_4_-DIBMA/DMPC nanodiscs at m_P_/m_L_ of 4. Insert shows 2D class averages of auto-picked particles from a total of 18 micrographs (Scale bar: 10 nm). (c) UV absorption spectra of mPEG_4_-DIBMA, DIBMA, and SMA(2:1). (d) Colloidal stability of mPEG_4_-DIBMA/DMPC nanodiscs at m_P_/m_L_ of 4 as determined by turbidity in response to increasing concentration of Mg^2^ or Ca^2+^. (e) Differential scanning calorimetry (DSC) thermograms displaying the excess molar isobaric heat capacities (ΔC_p_) of 4 mg/mL DMPC LUVs and mPEG_4_-DIBMA/DMPC nanodiscs at the indicated polymer/DMPC mass ratios, m_P_/m_L_. (f) Main phase transition temperature (T_m_) as a function of the polymer/DMPC mass ratio, m_P_/m_L_.

Next, we assessed the morphology of mPEG_4_-DIBMA/DMPC nanodiscs with the aid of negative-stain transmission electron microscopy (TEM). TEM micrographs and 2D classifications of auto-picked particles demonstrated the presence of homogeneously sized nanodiscs with an average diameter of ∼10 nm (Fig. 3b), consistent with our DLS data (Fig. 1). Furthermore, absorbance measurements showed that mPEG_4_-DIBMA has a very low absorbance in the ultraviolet (UV) spectral range, offering favorable properties for the photometric quantification of protein levels (Fig. 3c). In order to evaluate the effects of divalent cations on the colloidal stability of nanodiscs, we measured the turbidity of mPEG_4_-DIBMA/DMPC nanodiscs as a function of increasing concentrations of Mg^2+^ or Ca^2+^. mPEG_4_-DIBMA/DMPC nanodiscs began to precipitate at divalent ion concentrations of 10 mM Mg^2+^ or 5 mM Ca^2+^ (Fig. 3d and S3). Thus, mPEG_4_-DIBMA/DMPC nanodiscs showed a rather modest colloidal stability in the presence of divalent cations, as expected for negatively charged nanodiscs. Nevertheless, this modest stability is still sufficient for physiological concentrations of divalent cations and, thus, should enable activity studies of membrane proteins requiring typical concentrations of divalent cations.

To investigate the integrity of the nanodisc’s lipid matrix, we measured the effect of mPEG_4_-DIBMA on the thermotropic behavior of the encapsulated lipid-bilayer patch using differential scanning calorimetry (DSC). DSC thermograms showed that the phase-transition peak became broader with increasing *m*_P_*/m*_L_ (Fig. 3e). This observation confirms the formation of smaller lipid-bilayer nanoparticles upon addition of mPEG_4_-DIBMA to DMPC vesicles, which showed the expected highly cooperative gel-to-fluid transition at ∼24.5 °C (Fig. 3e, grey). It is worth noting that multiple phase transition peaks were observed at low polymer/lipid mass ratios, where nanodiscs coexist with vesicles (Fig. 3e and S4). Hence, it appears that a polymer/lipid mass ratio of ∼1 is required for mPEG_4_-DIBMA to fully solubilize DMPC vesicles, in agreement with our DLS data (Fig. 1c) and in contrast with SMA(2:1) (Fig. 3e and S4a). At higher polymer/lipid ratios, the mPEG_4_-DIBMA nanodiscs revealed a moderate decrease in the phase transition temperature (*T*_m_) to about 22 °C. We infer that the DMPC molecules along the perimeter of the nanodiscs were affected by mPEG_4_-DIBMA, whereas the core of the DMPC bilayer in the nanodiscs was not significantly affected. Similar results were observed in DSC measurement of DIBMA/DMPC nanodiscs (Fig. 3f and S4b), in line with previous findings.^27^ In contrast, SMA-encapsulated nanodiscs showed a steep drop in *T*_m_ with increasing polymer concentration down to 11.6 °C at *m*_P_*/m*_L_ of 4 (Fig. 3f and S4a). This observation has been explained by a perturbation of the lipids’ acyl chain packing by the polymer, which might be caused by the intercalation of the phenyl groups of SMA(2:1).^38^ Overall, our DSC data demonstrate that, in comparison with SMA(2:1), mPEG_4_-DIBMA requires slightly elevated polymer/lipid mass ratios for complete solubilization. However, mPEG_4_-DIBMA also has much milder effects on the lipid matrix under the conditions required to prepare small and homogenous nanodiscs, making it an ideal tool for subsequent biophysical or structural studies.

### Extraction capabilities of GPCRs from insect cell membranes

In addition to the solubilization of synthetic lipids, we tested the capacities of all new polymers to extract and encapsulate integral transmembrane proteins from cellular membranes into nanodiscs. For this purpose, we selected a prototypical GPCR comprising a thermostabilized mutant of the melanocortin 4 receptor (MC4R-eYFP) carrying an enhanced yellow fluorescent protein (eYFP) tag. We exposed crude membrane preparations from Sf9 insect cells to different concentrations of polymers and quantified the amounts of extracted MC4R by measuring the emission intensities of eYFP (Fig. 4). Interestingly, all four new DIBMA variants considerably enhanced the solubilization efficiency as compared with unmodified DIBMA (Fig. 4). Previous findings have shown that DIBMA carries higher negative charge densities than SMA(2:1) under similar conditions.^28^ As a result, strong Coulombic repulsion may affect the interaction of DIBMA with the negatively charged cell membrane and hinder the extraction of membrane proteins into native nanodiscs.

**Figure 4.**
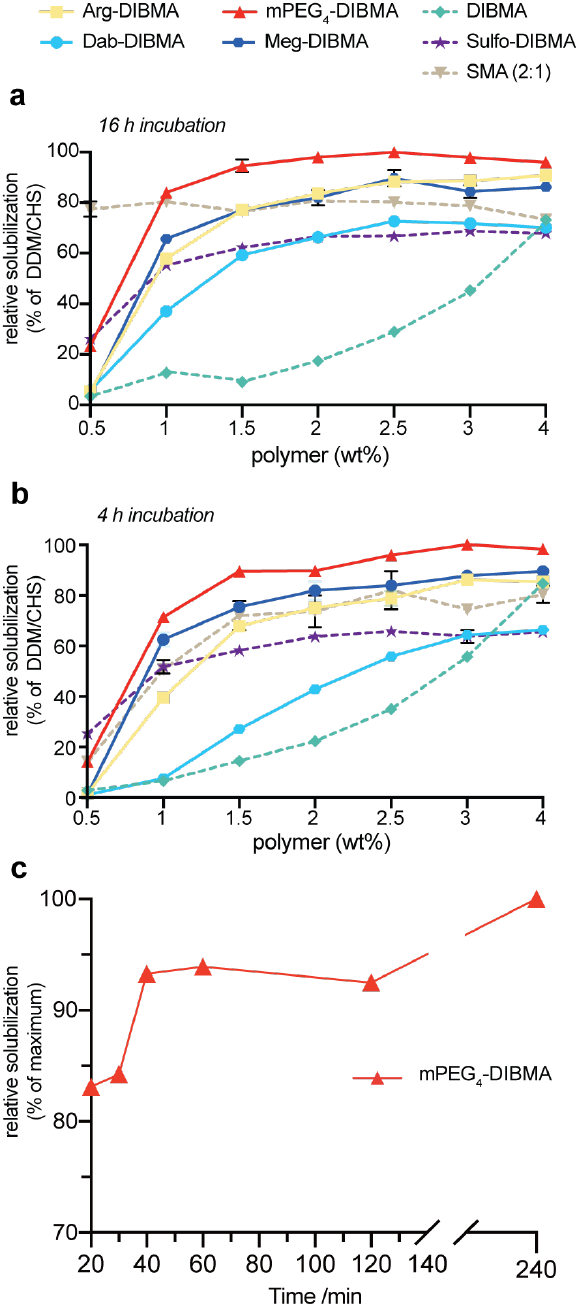
All new DIBMA variants, in particular mPEG_4_-DIBMA, efficiently extract the GPCR MC4R. (a,b) Polymer-mediated extraction of the melanocortin-4 receptor (MC4R-eYFP) from insect membranes. The extraction efficiencies of the different polymers in comparison with the frequently used DDM/CHS system (1% (w/v) DDM and 0.2% (w/v) cholesteryl hemisuccinate (CHS)) are plotted. The amount of solubilized receptor for each preparation was determined by eYFP fluorescence after incubation with the indicated polymers for either 16 h (a) or 4 h (b). (c) Normalized solubilization efficiency of 2% (w/v) mPEG_4_-DIBMA as a function of incubation time

Our data show that the newly designed DIBMA variants indeed facilitate the cell-membrane fragmentation process, resulting in higher extraction efficiencies. With the exception of Dab-DIBMA, all new DIBMA variants also offered solubilization efficiencies comparable to or surpassing that of SMA(2:1) at polymer concentrations ≥ 1%. Again, mPEG_4_-DIBMA displayed the best performance, approaching solubilization yields similar to the DDM/CHS system that was used as reference and generally serves as a gold standard for the extraction of GPCRs. In direct comparison with electroneutral Sulfo-DIBMA, the solubilization of MC4R was about 1.5-fold higher with mPEG_4_-DIBMA, making this new DIBMA variant particularly interesting for studies where the amount of extracted membrane protein is a limiting factor. Noteworthily, many functional and structural studies of GPCRs suffer from limited stability of these labile receptors. Therefore, long incubation times of the cell pellet with the polymer may adversely affect subsequent *in vitro* studies. To address this important point, we lastly tested the effects of reducing the incubation time (Fig. 4b, 4 h). These data largely reproduced the results obtained at longer incubation times (Fig. 4a, 16 h). The only exceptions were SMA(2:1) and Dab-DIBMA at low polymer concentrations, which displayed significantly lower extraction efficiencies at 4 h than at 16 h incubation. A more thorough analysis carried out for mPEG_4_-DIBMA further suggested that even shorter incubation times, in the range of 20–60 min, can be used to efficiently extract MC4R (Fig. 4c).

Taken together, we found that all new DIBMA variants can solubilize synthetic lipids and extract a prototypical GPCR from insect cells into native nanodiscs. Our MDS assay identified mPEG_4_-DIBMA as a new member of the small set of polymers that avert unspecific Columbic interactions with a cationic peptide ligand, thus facilitating interaction studies using nanodisc-embedded membrane proteins. Furthermore, we observed an outstanding performance of mPEG_4_-DIBMA in efficiently extracting MC4R from native cellular membranes. Paired with its decent Mg^2+^ and Ca^2+^ tolerance and its low UV absorbance, mPEG_4_-DIBMA has a particularly high potential for future applications in membrane-protein research.

## Conclusions

Polymer-encapsulated nanodiscs offer several advantages over other membrane mimetics, including the detergent-free extraction of target membrane proteins within their native lipid-bilayer environment. Several amphiphilic copolymers, such as SMA and DIBMA, have been successfully used for purifying membrane proteins from different cells and for determining membrane-protein structures.^31,39–41^ In general, existing polymers differ in key features such as extraction efficiency, tolerance against divalent metal ions, UV absorbance, and charge density, a factor that often restricts interaction studies using polymer-encapsulated nanodiscs. Although all of the listed properties are essential for characterizing membrane proteins such as GPCRs, none of the currently available polymers combine all desirable properties, necessitating compromises to be made in the selection process. To overcome this limitation, we here introduced and tested a series of new DIBMA variants. All new polymers show promising features, in particular as regards their solubilization efficiency of a GPCR from insect membranes. A PEGylated polymer, mPEG_4_-DIBMA, emerged as the top performer in all tested properties. mPEG_4_-DIBMA overcomes some of the most serious limitations of SMA and DIBMA without sacrificing protein yields, highlighting its potential for the functional and structural characterization of membrane proteins in their native lipid-bilayer environment.

## Author Contributions

CC carried out biophysical and cell biology experiments; SS recorded and analyses EM data; CV and SK designed polymers, CV and MCB synthesized polymers, ME, SK and GFS designed, supervised, and discussed experiments; CC, CV, SK and ME wrote the manuscript. All authors contributed to data analysis, scientific discussions and content of the manuscript.

## Acknowledgements

We thank Prof. Dr. Georg Groth (Heinrich Heine University Düsseldorf) for granting access to a DLS instrument, Prof. Dr. Raymond C. Stevens (iHuman Institute at ShanghaiTech University) for providing us with a plasmid encoding the thermostabilized melanocortin 4 receptor, and Sarah Tutz (University of Graz) for excellent technical assistance. This work was supported by the German Research Foundation (DFG) (103/4-1, ET 103/4-3, and the Heisenberg grant ET 103/5-1) to M.E., the CSC (No. 201708320246) to C.C., and the Austrian Science Fund FWF (project I 5359-N) to S.K

