## Supplementary Information for "Capturing G protein-coupled receptors into native lipid-bilayer nanodiscs using new diisobutylene/maleic acid (DIBMA) copolymers"

### 1. Supplementary Procedures

**Materials.** The zwitterionic, saturated phospholipid 1,2-dimyristoyl-*sn*-glycero-3-phosphocholine (DMPC) was obtained from Avanti Polar Lipids (Avanti, Alabaster, USA). DIBMA monomethyl ester was a kind gift from Glycon Biochemicals (Luckenwalde, Germany). Styrene/maleic acid (SMA(2:1)) copolymer was purchased from Orbiscope (Geleen, The Netherlands). *n*-Dodecyl-β-D-maltopyranoside (DDM) was from Glycon Biochemicals (Luckenwalde, Germany). L-Arginine was purchased from Carl Roth (Karlsruhe, Germany), meglumine from Fisher Scientific (Schwerte, Austria), and L-2,4-diaminobutyric acid dihydrochloride from Bachem (Bubendorf, Switzerland). $MgCl_2$, $CaCl_2$, cholesteryl hemisuccinate (CHS), tris(hydroxymethyl)aminomethane hydrochloride (Tris-HCl), 2-(4-(2-hydroxyethyl)-1-piperazinyl)-ethanesulfonic acid (HEPES), and other chemicals were purchased from Sigma–Aldrich (Darmstadt, Germany).

**Syntheses.** All starting materials were purchased from TCI (Eschborn, Germany), Fisher Scientific (Schwerte, Austria), Th. Geyer (Renningen, Germany), and Sigma–Aldrich (Darmstadt, Germany) and were used as received. The synthetic procedures for DIBMA amide derivatives followed the same general protocol previously published for Glyco-DIBMA.[1] Briefly, to DIBMA monomethyl ester (2 g, 15.6 mmol) dissolved in 80 mL MeOH was added an amine (15.6 mmol; see below) in 25 wt% sodium methoxide (4 mL diluted in 20 mL MeOH) under stirring at room temperature. The reaction mixture was refluxed overnight, and MeOH was removed by rotary evaporation. The identity of the resulting DIBMA amide product was confirmed by attenuated total reflection infrared (ATR-IR) spectroscopy. Arg-DIBMA: L-arginine was used as amine; ATR-IR: 2942 (C–H str), 1722 (C=O str), 1558 (C–N str/NH bend) $cm^{-1}$; Dab-DIBMA: L-2,4-diaminobutyric acid dihydrochloride was used as amine; ATR-IR: 3345, 3283, 3178 (N–H str), 2946 (C–H str), 1727 (C=O str), 1567 (C–N str/ NH bend) $cm^{-1}$; Meg-DIBMA: meglumine was used as amine; ATR-IR: 3296 (O–H str), 2943 (C–H str), 1726 (C=O str), 1575 (C–N str/NH bend) $cm^{-1}$; $mPEG_4$-DIBMA: tetraethyleneglycol monomethyl ether amine ($mPEG_4$-amine) was used as amine; ATR-IR: 2868 (C–H str), 1729 (C=O str), 1582 (C–N str/ NH bend) $cm^{-1}$.

The general synthetic procedure for preparing $mPEG_4$-amine [2–4] is shown below:

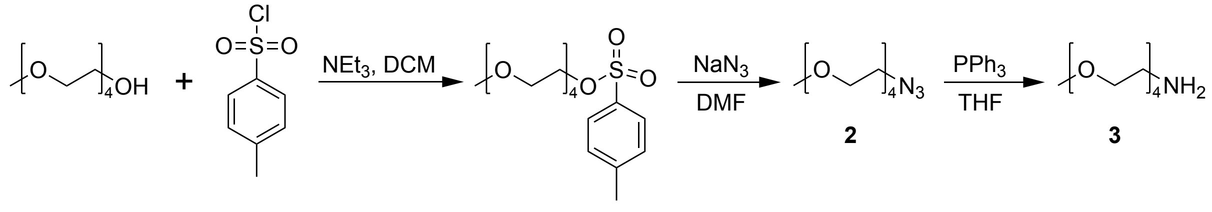


1 2 3

2,5,8,11-Tetraoxatridecan-13-yl 4-methylbenzenesulfonate (**1**)

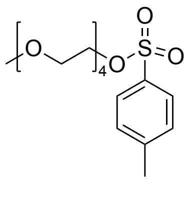

A solution of tetraethylene glycol monomethyl ether (10.0 g, 48 mmol, 1 eq.) in DCM (50 mL) was added dropwise over 30 min to a solution of *p*-toluenesulfonyl chloride (9.16 g, 48 mmol, 1 eq.) suspended in DCM (50 mL) while stirring at room temperature. Triethylamine (13.4 mL, 0.096 mmol, 2 eq.) was then added dropwise over 10 min and the reaction mixture was stirred at room temperature for 22 h. The solvent was evaporated under reduced pressure and the residue was diluted in DCM (70 mL) and washed with NaOH (1 M, 3 × 70 mL), followed by water (3 × 70 mL). The organic phase was dried over $MgSO_4$, filtered, and concentrated under reduced pressure. The resulting oil was purified by gravity column chromatography using ethyl acetate as eluent. The product was obtained as a yellowish oil in 73% yield. The identity of the resulting product was confirmed by NMR and attenuated total reflection infrared (ATR-IR) spectroscopy: $^{1}$H-NMR (300 MHz, $CDCl_3$, 25°C): $\delta$ (ppm) = 2.39 (s, 3H, Ar–$CH_3$), 3.32 (s, 3 H, –O$CH_3$), 3.45–3.66 (m, 14 H, 3(–O–$CH_2CH_2$–O–), –O$CH_2$–), 4.10 (t, 2H, Ts–$CH_2$–, $^{3}J_{H\text{–}H}$ = 4.89 Hz), 7.29 (d, 2H, Ar–H, $^{3}J_{H\text{–}H}$ = 8.01 Hz), 7.74 (d, 2H, Ar–H, $^{3}J_{H\text{–}H}$ = 8.32 Hz); $^{13}$C-NMR (75 MHz, $CDCl_3$, 25°C): $\delta$ (ppm) = 144.9, 133.1, 129.9, 128.1, 72.0, 70.8, 70.7, 70.6, 70.6, 70.6, 69.4, 68.8, 59.1, 21.7; ATR-IR[2,3] ($cm^{-1}$): 2874 (C–H str, aliphatic), 1452 (C=C str, aromatic), 1353 (S=O str, asym), 1189 (S=O str, sym), 1175 (C–C str, aliphatic), 1096 (C–O–C str).

13-Azido-2,5,8,11-tetraoxatridecane (**2**)

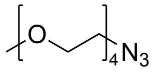

To a suspension of sodium azide (1.54 g, 23.4 mmol, 1.5 eq.) in DMF (40 mL), a solution of **1** (5.66 g, 15.6 mmol, 1.0 eq.) in DMF (25 mL) was added dropwise over 50 min and the reaction mixture was stirred at room temperature for 22 h. The mixture was diluted with water (130 mL) and washed with DCM (7 × 100 mL). The organic fractions were collected and washed further with water (2 × 100 mL), dried over $MgSO_4$, filtered, and concentrated under reduced pressure. DMF was removed by repeated distillation of the solution with toluene. The residue was purified by gravity column chromatography using 2:1 hexane/EtOAc as eluent. The product was obtained as a colorless oil in 86% yield. The identity of the resulting product was confirmed by NMR and attenuated total reflection infrared (ATR-IR) spectroscopy: $^{1}$H-NMR (300 MHz, $CDCl_3$, 25°C): $\delta$ (ppm) = 3.27–3.41 (m, 5H, –O$CH_3$, –$OCH_2$–$CH_2$–$N_3$), 3.48–3.55 (m, 2H, –O$CH_2$–$CH_2$–$N_3$, 3.57–3.68 (m, 12H, 3 × –O$CH_2$–$CH_2$O–); $^{13}$C-NMR (75

MHz, $CDCl_3$, 25°C): $\delta$ (ppm) = 72.1, 70.8, 70.8, 70.8, 70.8, 70.7, 70.2, 59.2, 50.8; ATR-IR[2,3] ($cm^{-1}$): 2870 (C–H str, aliphatic), 2098 (C–$N_3$), 1101 (C–O–C str).

2,5,8,11-Tetraoxatridecan-13-amine (**3**)

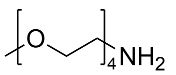

To a solution of **2** (3.0 g, 12.9 mmol, 1.0 eq.) in anhydrous THF (25 mL), triphenylphosphine (4.22 g, 16.1 mmol, 1.25 eq.) in anhydrous THF (15 mL) was added dropwise over 15 min under nitrogen atmosphere and the reaction mixture was stirred at room temperature for 5 h. Distilled water (20 mL) was then added and the solution was refluxed for 22 h. The solution was concentrated under reduced pressure and the residue was purified by gravity column chromatography using 9:1 DCM/MeOH (+1% $Et_3N$). The product was obtained as a yellowish oil in 97% yield. The identity of the resulting product was confirmed by NMR and attenuated total reflection infrared (ATR-IR) spectroscopy: $^1$H-NMR (300 MHz, $CDCl_3$, 25°C): $\delta$ (ppm) = 2.82 (t, 2H, $^3J_{H\text{–}H}$ = 5.21 Hz, –$CH_2$–$CH_2$–$NH_2$), 3.33 (s, 3H, –O$CH_3$), 3.43–3.54 (m, 4H, –O$CH_2$–$CH_2$O–, –$CH_2$–$CH_2$–$NH_2$), 3.54–3.68 (m, 10H, 2× –O$CH_2$–$CH_2$O, –O$CH_2$–$CH_2$O–); $^{13}$C-NMR (75 MHz, $CDCl_3$, 25°C): $\delta$ (ppm) = 73.3, 72.1, 70.7, 70.7 (2×), 70.6, 70.4, 59.2, 41.8; ATR-IR[2,3] ($cm^{-1}$): 3372 (N–H str), 2866 (C–H str, aliphatic), 1100 (C–O–C str).

**Polymer stock solutions.** Polymer powders used throughout this study were dissolved in 50 mM Tris-HCl or 50 mM HEPES, 200 mM NaCl unless noted otherwise, followed by incubation at 55 °C for 10–15 min with vortexing in an alternating fashion to yield a mass concentration of 50 mg/mL. The final pH value of polymer stock solutions was adjusted to 8.0 or 7.4, as indicated. All polymer stock solutions were then filtered through polycarbonate membranes with a pore size of 0.22 µm and stored at –4 °C.

**Large unilamellar vesicles (LUVs).** DMPC powder was dissolved in 50 mM Tris-HCl or HEPES, 200 mM NaCl, pH 8.0 to a final concentration of 20 mg/mL. The lipid suspension was incubated at 35 °C for 30–40 min with vortexing every 10 min in an alternating fashion. In order to increase the hydration efficiency, the lipid suspension was then immersed in liquid nitrogen followed by thawing in a ThermoMixer (Eppendorf, Germany) at 35 °C for 5–10 freeze–thaw cycles. Subsequently, the DMPC suspension was loaded into gas-tight syringes and extruded at 35 °C through a 100-nm Whatman polycarbonate membrane filter at least 20 times using a Mini-Extruder (Avanti, Alabaster, USA) to prepare LUVs.

**Ultraviolet–visible (UV–vis) spectroscopy.** The absorption spectra of polymers at 1 mg/mL dissolved in 50 mM Tris-HCl, 200 mM NaCl, pH 8.0 were recorded on a Jasco V-650 UV–vis spectrophotometer

(JASCO Germany). Measurements were performed at room temperature using a quartz cuvette with a 10-mm light path (Hellma Analytics, Germany) at a scan rate of 200 nm/min in the wavelength range of 220–600 nm.

**Polymer/lipid nanodiscs.** The DMPC LUVs suspension was added to the polymer stock solutions at different polymer/lipid mass ratios ($m_P/m_L$) and incubated at 35 °C for at least 16 h with shaking at 450 rpm to form polymer-encapsulated DMPC nanodiscs. All experiments were performed in aqueous buffer containing 50 mM Tris-HCl, 200 mM NaCl, pH 8.0.

**Dynamic light scattering (DLS).** Solubilization efficiencies of all polymers were confirmed and quantified with the aid of DLS using DMPC model membranes. A Zetasizer Nano S (Malvern Panalytical, UK) was used to perform DLS measurements in a 70-μL microcuvette (Brand, Wertheim, Germany) at a temperature of 25 °C. The DLS instrument was equipped with a He–Ne laser having a wavelength of 633 nm, and the detection scattering angle was 173°. The sample was thermostatted for 5 min at 35 °C prior to measurement.

**Differential scanning calorimetry (DSC).** DSC measurements were carried out using a MicroCal VP-DSC calorimeter (Malvern Panalytical, UK) to monitor the thermotropic behavior of DMPC lipids in the absence and presence of polymers. Samples were prepared in 50 mM Tris-HCl, 200 mM NaCl, pH 8.0 at various polymer/lipid mass ratios ($m_P/m_L$). Polymer/DMPC samples and reference buffer were first degassed and then measured in 5–10 heating and cooling cycles at a scan rate of 60 °C $h^{-1}$ in the range of 2–80 °C. One representative curve was chosen from the closely overlaid scan traces, the buffer baseline was subtracted, and the data were normalized against the DMPC concentration (6 mM) using MicroCal Origin 7.0 software (OriginLab, Northampton, USA). The phase transition temperature ($T_m$) was determined as the temperature at which the excess molar isobaric heat capacity ($C_p$) reached its maximum value. The size of the cooperative unit (CU) was obtained as the ratio of the van't Hoff enthalpy, given by the shape of the melting peak, to the calorimetric enthalpy, given by the peak area.[5]

**Turbidity experiments.** The colloidal stability of mPEG$_4$-DIBMA lipid particles in the presence of divalent cations was evaluated with turbidity experiments at 620 nm using a Tecan Spark 10M microplate reader (Tecan, Switzerland).

**ζ-potential measurements.** Samples containing 4 mg/mL DMPC LUVs, 8 mg/mL polymer, or polymer/DMPC nanodiscs at $m_P/m_L$ = 2 were prepared in 50 mM Tris-HCl, 100 mM NaCl, pH 7.4. ζ-potential measurements were carried out on a Zetasizer Nano ZS (Malvern Panalytical, UK) using disposable folded capillary zeta cells DTS1070 (Malvern Panalytical, UK) at a detection scattering angle of 173° and a temperature of 25 °C. The diffusion barrier technique was used to prepare the sample cell for $\zeta$-potential measurement.[6] The folded capillary cell was first filled with buffer before 100 μL of sample was pipetted directly into the cell bottom with the aid of a gel-loading tip, avoiding mixing of the sample with buffer during loading. The cell was capped and equilibrated for 300 s prior to measurement to reduce fluid motion induced by temperature gradients.

**Adrenocorticotropic hormone peptide (ACTH).** The human adrenocorticotropic hormone truncated construct ACTH (1-23)Cys was subcloned into pET-16b vector, including a 6xHis tag at N-terminus, followed by B1 domain of streptococcal protein G (GB1 fusion protein) and tobacco etch virus (TEV) cleavage site. *Escherichia coli* strain BL21 (DE3) cells were used for peptide expression. Cells were cultured in 2xYT medium at 37 °C, 140 rpm in baffled flask, supplemented with 100 μg/mL ampicillin. The expression was inducted with 1 mM isopropyl-β-D-thiogalactopyranoside (IPTG) upon the optical density 600 ($OD_{600}$) reached 0.6–0.8. Cells were harvested by centrifugation at 6,000 *g*, 10 min, 4 °C after 5 h induction, flash-frozen in $N_2$ (l), and stored at –80 °C for future use. Cell pellets were resuspended in lysis buffer containing 50 mM sodium phosphate, pH 7.5, 300 mM NaCl, 1 mM dithiothreitol (DTT), supplemented with EDTA free cOmplete protease inhibitor cocktail tablet (Roche). The homogenate was sonicated on ice, and the lysate was centrifuged at 100,000 *g* for 25 min at 4 °C. The supernatant was incubated with Ni-NTA resin (Macherey-Nagel, Germany) at 4 °C overnight with gentle rotation. The resin was loaded onto a gravity column and washed with 10 column volumes (CVs) lysis buffer and 10 CVs lysis buffer supplemented with 20 mM imidazole. The peptide was eluted with 10 CVs lysis buffer supplemented with 250 mM imidazole. Fractions containing peptides were collected and dialyzed against lysis buffer using 3.5K MWCO SnakeSkin dialysis tubing (ThermoFisher) at 4 °C overnight. The fusion protein was removed by addition of home-made TEV protease. The cleaved peptide was further purified by size-exclusion chromatography (Superdex 16/600, 30 pg column; GE health) with a running buffer containing 20 mM ammonium bicarbonate. Peptide labeling with thiol-reactive ATTO-488 maleimide (ATTO-TEC, Germany) was carried out according to the manufacturer's instructions. The labeled peptide was further purified using high-performance liquid chromatography (HPLC) using a Zorbax SB300 C8 4.6 x 250 mm analytical column (Agilent Technologies).

**Microfluidic diffusional sizing (MDS).** ATTO 488-labeled human adrenocorticotropic hormone (ACTH) peptide was added to polymer/DMPC nanodiscs at $m_P/m_L = 4$ to a final peptide concentration of 20 nM. The final concentration of DMPC was 4 mg/mL. Interactions between peptide and polymer/DMPC nanodiscs were measured on a Fluidity One-W system (Fluidic Analytics, Cambridge, UK) by recording changes in hydrodynamic particle size.[7,8] Disposable MDS chips were used holding a total sample volume of 5 μL. All experiments were carried out in 50 mM HEPES, 200 mM NaCl, pH 7.4 or 8.0.

**Negative-stain transmission electron microscopy (TEM).** TEM specimens were prepared by spreading 4 µL mPEG$_4$-DIBMA/DMPC nanodiscs at $m_P/m_L = 4$ onto freshly glow-discharged copper grids (15 mA for 25 s at 0.39 mbar) coated with a carbon support film. Excess suspension was blotted off after ~5 s using filter paper. Grids were washed with 5 µL MilliQ water once, followed by staining with 5 µL 2% (*w*/*v*) aqueous uranyl acetate solution twice and blotted off after ~15 s with filter paper. After preparation, specimens were air-dried and examined on a Talos L120C transmission electron microscopy equipped with a 4k × 4K Ceta CMOS camera (Thermo Scientific).

**Preparation of insect membranes and solubilization of MC4R.** The wild-type human melanocortin 4 receptor (MC4R, UniProtKB-P32245) was codon-optimized and inserted into a mortified pFastbac1 vector, including an influenza virus hemagglutinin (HA) signal peptide followed by a Flag tag at the N-terminus as well as a human rhinovirus (HRV3C) cleavage site and a 10xHis tag at the C-terminus by using NcoI-EcoRI restriction endonucleases (New England Biolabs). The thermostabilized mutant MC4R construct was modified by introducing 5 mutations (E49$^{1.37}$V, N97$^{2.57}$L, S99$^{2.59}$F, S131$^{3.34}$A and D298$^{7.49}$N, where superscript numbers correspond to the Ballesteros–Weinstein nomenclature[9]) containing the same tags as the wild type, a generous gift of Prof. Dr. Raymond C. Stevens (iHuman Institute at ShanghaiTech University).[10] The MC4R-eYFP construct was modified by introducing the eYFP gene between the HRV3C cleavage site and the 10xHis tag. Recombinant baculovirus production of MC4R-eYFP was generated by transfecting *Spodoptera frugiperda* (Sf9) cells grown in Sf-900 III SFM media (ThermoFisher) at 27 °C with Bacmid (Bac-to-Bac system, ThermoFisher) using FuGENE HD transfection reagent according to the manufacturer's instructions. Sf9 cells were infected at a density of 2 to 3 x $10^6$ cells/mL with recombinant baculovirus. 72 h after infection, cells were harvested by centrifugation at 6000 *g*, 4 °C for 10 min and stored at –80 °C. Cellular membranes were prepared as described previously with slight modifications.[10] In brief, Sf9 cell pellets were resuspended in a hypotonic buffer containing 10 mM HEPES, 10 mM $MgCl_2$, 20 mM KCl, pH 7.8 and cOmplete EDTA-free protease inhibitor cocktail tablet (Roche) and then lysed by sonication on ice (Bandelin Sonorex, Germany). Cell membranes containing MC4R-eYFP were harvested by ultracentrifugation at 120,000 *g*, 4 °C for 30 min (Optima xpn-80 ultracentrifuge, Beckman Coulter, USA). The membrane pellet was resuspended in the same hypotonic buffer and disrupted with a Dounce homogenizer followed by ultracentrifugation at 120,000 *g*, 4 °C for 30 min. The above procedure was repeated twice. Further membrane purification was performed three times using hypotonic buffer supplemented with 1 M NaCl. Purified membrane pellets were washed once with 50 mM HEPES, 200 mM NaCl, pH 8.0, flash-frozen in liquid nitrogen, and stored at –80 °C. Polymers were added to 20 mg/mL (wet mass/volume) purified cellular membranes at different concentrations and incubated at 4 °C for either 4 h or 16 h with gentle rotation. Supernatants were collected after ultracentrifugation at 120,000 *g*, 4 °C for 30 min.

#### 2. Supplementary Figures

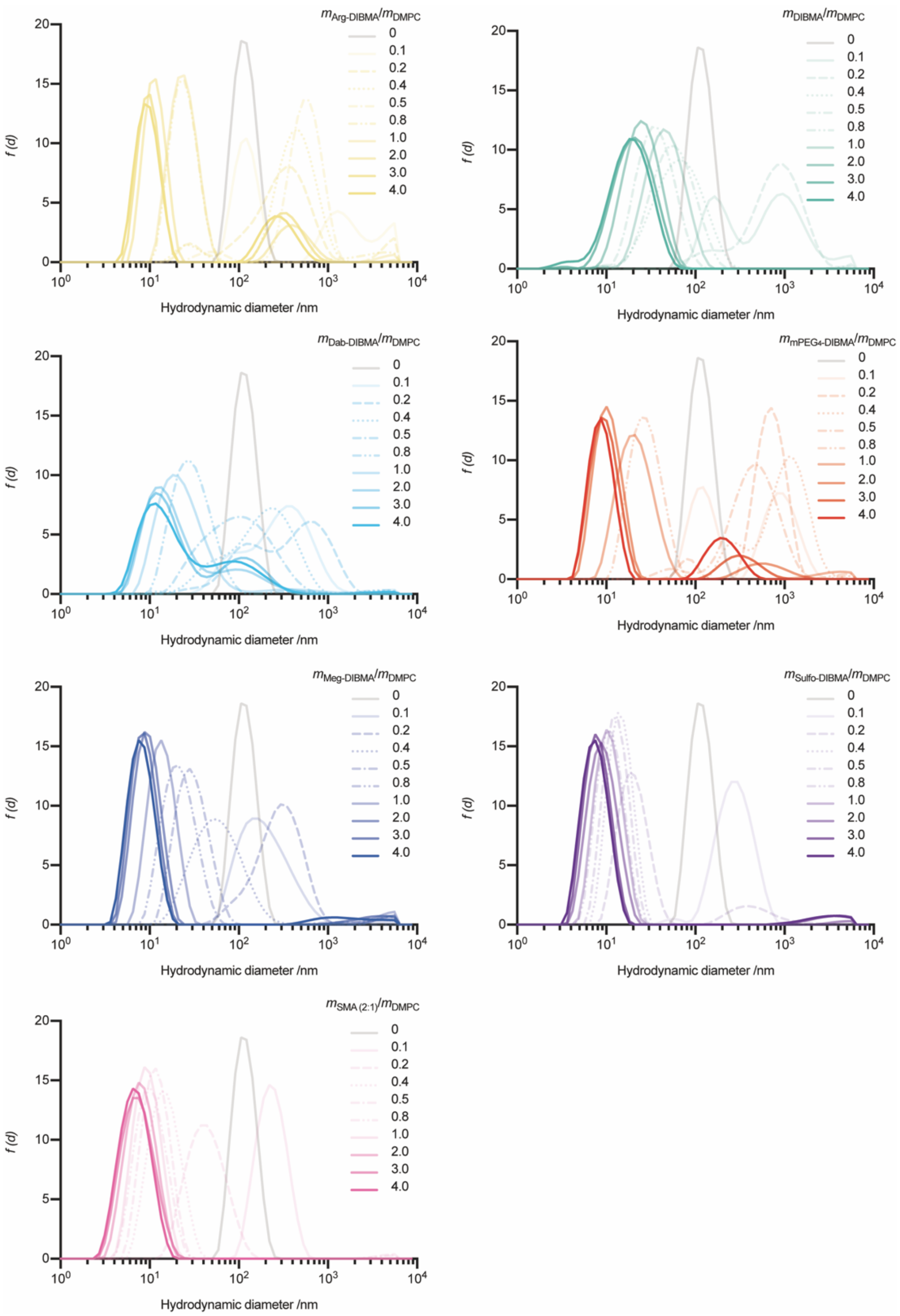


**Supplementary Figure S1.** Solubilization of DMPC vesicles and formation of polymer-encapsulated DMPC nanodiscs. Intensity-weighted particle size distributions of mixtures containing polymer and DMPC (4 mg/mL) at indicated polymer/lipid mass ratios ($m_P/m_L$).

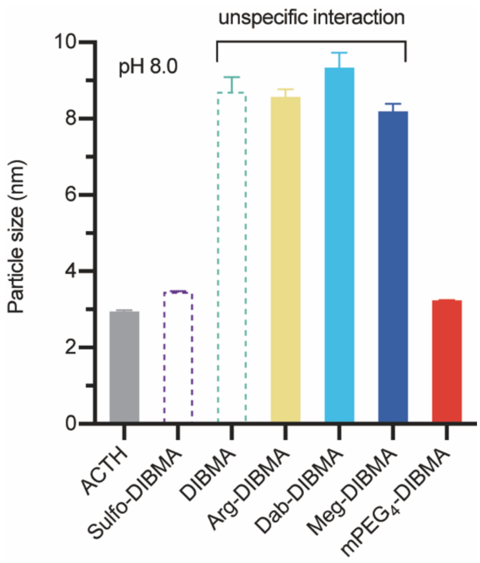


**Supplementary Figure S2.** Unspecific interactions between peptide and polymer/DMPC nanodiscs at $m_P/m_L = 4$ were measured by means of microfluidic diffusional sizing (MDS). All experiments were carried out at 50 mM HEPES, 200 mM NaCl, pH 8.0. Error bars indicate ± one standard deviation of two independent experiments, each repeated in triplicate.

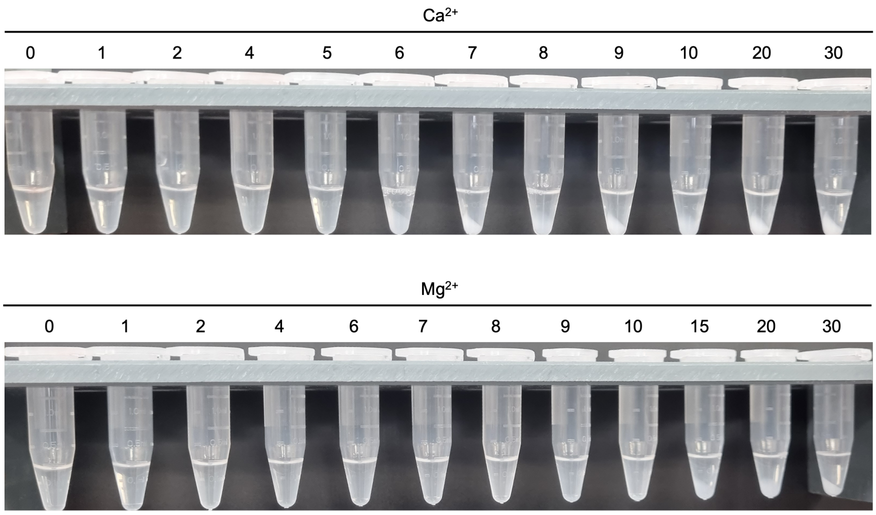


**Supplementary Figure S3.** Visual appearance of mPEG4-DIBMA/DMPC nanodiscs at $m_P/m_L = 4$ in response to increasing concentration of $Mg^{2+}$ or $Ca^{2+}$.

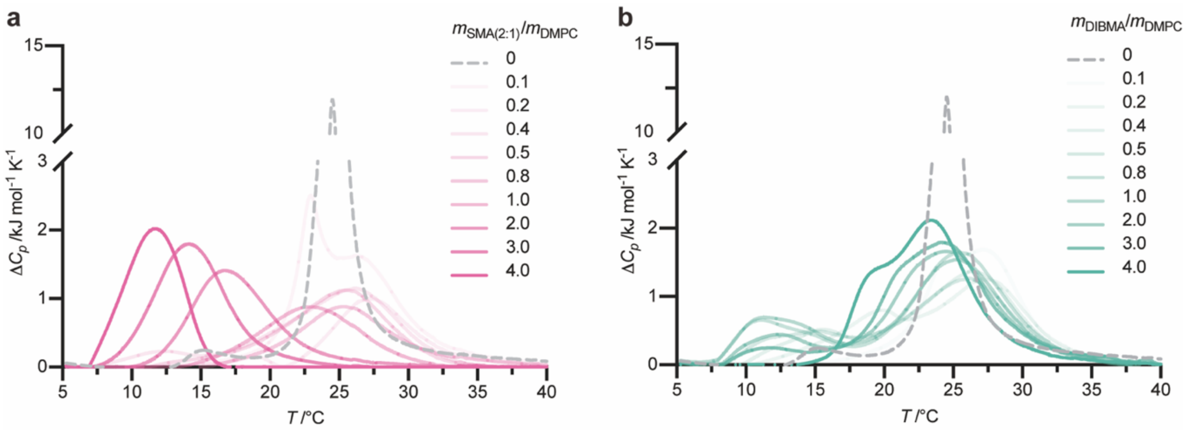


**Supplementary Figure S4.** DSC thermograms of polymer-encapsulated DMPC nanodiscs. DSC thermograms displaying excess molar isobaric heat capacities ($\Delta C_p$) of SMA(2:1)/DMPC (a) and DIBMA/DMPC (b) at indicated polymer/DMPC mass ratios, $m_P/m_L$.
